# CircSMAD3 represses VSMC phenotype switching and neointima formation via promoting hnRNPA1 ubiquitination degradation

**DOI:** 10.1101/2023.09.17.558158

**Authors:** Shuai Mei, Li Zhou, Xiaozhu Ma, Qidamugai Wuyun, Chen Chen, Ziyang Cai, Jiangtao Yan, Hu Ding

## Abstract

**Aims:** Circular RNAs (circRNAs) are novel regulatory RNAs with high evolutionary conservation and stability, making them attractive therapeutic agents for various vascular diseases. SMAD family is a downstream mediator of the canonical TGFβ signalling pathway and has been considered a critical regulator in vascular injury. However, the role of circRNAs derived from the SMAD family members in vascular physiology remains unclear. Therefore, this study aimed to identify a functional circRNA derived from the SMAD family and elucidate its potential as an effective therapeutic agent for vascular proliferative diseases.

**Methods and Results:** We initially identified potential functional circRNAs originating from the SMAD family using integrated transcriptome screening. circSMAD3, derived from the SMAD3 gene, was identified to be significantly downregulated in vascular injury and atherosclerosis. Transcriptome analysis was conducted to comprehensively illustrate the pathways modulated by circRNAs. Functionally, circSMAD3 repressed VSMC proliferation and phenotype switching *in vitro* evidenced by morphological assays and ameliorated arterial injury-induced neointima formation *in vivo*. Mechanistically, circSMAD3 interacted with heterogeneous nuclear ribonucleoprotein A1 (hnRNPA1) within the nucleus, enhanced its interaction with E3 ligase WD repeat domain 76 (WDR76) to promote hnRNPA1 ubiquitination degradation, facilitated p53 pre-RNA splicing, activated p53γ signalling pathway, and finally suppressed VSMC proliferation and phenotype switching.

**Conclusion:** Our study identifies circSMAD3 as a novel epigenetic regulator that suppresses VSMC proliferation and phenotype switching, thereby attenuating vascular remodelling and providing a new circRNA-based therapeutic strategy for cardiovascular diseases.

## Introduction

Vascular smooth muscle cells (VSMCs) are the prominent cells in the blood vessel wall that can transition from a contractile state to a synthetic state in a process known as “phenotype switching” ^1, 2^, followed by the reduction of differentiation markers^3, 4^, such as α-smooth muscle actin (α-SMA), transgelin, and Calponin. When arterial injury occurs, this characteristic of VSMC leads to arterial lumen stenosis or structural abnormalities, which triggers the occurrence of vascular proliferative diseases, including atherosclerosis and restenosis after angioplasty and eventually increases the risk of myocardial infarction^5^. However, the underlying mechanisms of VSMC phenotype switching are partly understood.

As a novel type of non-coding RNA, circular RNAs (circRNAs) have been shown to significantly affect pathophysiological processes, including cardiovascular diseases^6, 7^. They act as microRNA sponges^8^, protein modulator^9, 10^, genetic transcription regulator^11^, and translation template^12, 13^ in pathophysiological processes, including circ_Lrp6^14^ and circEsyt2^15^ in vascular proliferative diseases. Because of their covalently closed-loop structures, CircRNAs are considered more stable and have recently attracted more attention for the recent development of synthetic circRNAs as theranostics and vaccines^16, 17^. To date, many attempts have been made to develop circRNA-based therapies and great progress has been made. For example, circRNAs have been successfully studied as vaccines for overcoming the coronavirus disease 2019 pandemic by engineering circRNAs that encode the severe acute respiratory syndrome coronavirus 2 receptor-binding domain^18^. These studies have revealed the great value and broad application prospects of circRNAs in cardiovascular diseases. Specifically, these results have shown that circRNAs have great potential as novel therapies for addressing cardiovascular diseases.

The SMAD family, as mediators of the canonical transforming growth factor-beta (TGF-β) signalling pathway, reportedly plays a critical role in vascular homeostasis^19–21^. Many studies have reported that mutations in the SMAD family members cause angiodysplasia or aortic aneurysm and dissection syndromes^22–24^. However, the role of circRNAs derived from the SMAD family members in vascular physiology remains unclear. Therefore, this study aimed to identify circSMAD3 to fully illustrate its vital functions in VSMC phenotype switching and proliferation *in vitro* and *in vivo* by enhancing the interaction of heterogeneous nuclear ribonucleoprotein A1 (hnRNPA1) with E3 ligase WD repeat domain 76 (WDR76), repressing the splicing of p53 precursor RNA (p53 pre-RNA), and facilitating p53γ signalling pathway. Our study demonstrates the critical role of circSMAD3 in vascular injury and provides novel targets for developing circRNA-based therapies for cardiovascular diseases.

## Methods

### Cell culture and treatment

Human primary aortic smooth muscle cell (HASMC) was purchased from Anwei Biotechnology Co., Ltd (Shanghai, China) and cultured in ICell Primary Smooth Muscle Cell Low Serum Culture System (Anwei Biotechnology Co, Shanghai, China) in a humidified atmosphere at 37°C with 5% CO2. Mouse primary aortic smooth muscle cells (mASMC) were isolated from adult mice aged 8 weeks and was cultured in Dulbecco’s Modified Eagle’s Medium (DMEM) containing 10% fetal bovine serum (FBS, GIBCO, Brazil) in a humidified atmosphere at 37°C with 5% CO2. Passages 4-15 of HASMC and mASMC were applied to this study.

### RNA isolation, quantitative RT-PCR and PCR

Total RNA was extracted using TRIzol reagent (Vazyme, Nanjing, China), as previously described. The purity and concentration of the total RNA were determined using a NanoDrop ND-2000 (NanoDrop Thermo, Wilmington, DE, USA). Next, the reverse transcription of circRNAs was performed using the Takara System (Dalian, China). Real-time polymerase chain reaction (PCR) was performed using the Vazyme System (Vazyme, Nanjing, China) on a 7900HT FAST Real-time PCR System (Life Technologies, Carlsbad, CA). The level of genes was calculated based on the cycle threshold (Ct) values compared to a reference gene using the formula 2^-ΔΔCt^. Glyceraldehyde-3-phosphate dehydrogenase (GAPDH) messenger RNA (mRNA) was used as a reference for mRNA and circRNA expression. PCR was performed according to the manufacturer’s instructions (NEO, Japan). The details of the primers used are listed in the Supplementary File.

### Protein extraction and western blotting

Briefly, the cells were lysed in ice-cold immunoprecipitation (IP) lysis buffer (Beyotime, Shanghai, China) and centrifuged at 12,000×g for 15 min at 4°C. The supernatant was collected, and the protein concentration was analysed using a NanoDrop ND-2000 (Thermo, Wilmington, DE, USA). Next, the protein levels were normalised by probing the same blots with antibodies against α-Tubulin (Abclonal, Wuhan, China) or GAPDH (Abclonal, Wuhan, China). The primary antibodies were anti-α-SMA antibody (1:200, Abclonal, China) and anti-hnRNPA1 antibody (1:200, Proteintech, China). The primary antibodies used in this study are listed in the Supplementary File.

### Fluorescence *in situ* hybridisation

The circSMAD3 probe labelled with cyanine 3 (Cy3) was designed and synthesised by RiboBio (Guangzhou, China), and the protocol was performed using a Fluorescent *in Situ* Hybridization Kit (RiboBio, Guangzhou, China). Furthermore, the probe signals were detected, and images were acquired using a Lei TCS SP8 Laser Scanning Confocal Microscope (Leica Microsystems, Mannheim, Germany).

### Immunofluorescence analysis

Briefly, HASMC and mASMC were washed with phosphate-buffered saline (PBS), fixed in 4% paraformaldehyde, permeabilised with 0.5% Triton X-100 (Beyotime, Shanghai, China) in PBS for 15 min, and incubated with 5% bovine serum albumin for 1 h at room temperature. Subsequently, the cells were incubated with primary antibodies (1:100) at 4°C overnight. The primary antibodies used included anti-α-SMA (1:200, Abclonal, China), anti-hnRNPA1 (1:200, Proteintech, China), and anti-WDR76 (1:200, Proteintech, China) antibodies. After the overnight incubation, the samples were incubated with secondary antibodies (FITC mouse, CY3 rabbit; Servicebio, Wuhan, China) at 1:500 for 1 h, washed three times with PBS, and then incubated with 4’,6-Diamidino-2-phenylindole (DAPI) for 15 min. The images were captured using a fluorescence microscope (Axio Imager 2, Zeiss, Germany).

### Cell area measure

First, HASMC and mASMC were seeded onto glass coverslips in 24-well plates. The cells were washed two times with PBS, fixed in 4% paraformaldehyde, and permeabilised with 0.5% Triton X-100 for 15 min. Subsequently, the samples were incubated with 0.1% phalloidin (Abclonal, Wuhan, China) for 30 min at 37°C, followed by incubating with DAPI at 1:100 dilution for 15 min. Coverslips were placed onto glass slides using a mounting medium, and the images were captured using a fluorescence microscope (Axio Imager 2, Zeiss, Germany). This process was repeated three times.

### Pull-down assay with biotinylated circSMAD3 probe

The biotinylated circSMAD3 probe was synthesised by AokeBoTai (Wuhan, China), and the oligo-probe was used as a control. This assay was performed as previously described. Briefly, the circSMAD3 probe was incubated with streptavidin magnetic beads (Life Technologies, USA) at room temperature for 1 h to generate probe-coated beads. The cell lysates were incubated with the probe-coated beads at room temperature for 2 h. Next, the complexes were washed two times and divided into two parts for RNA and protein extraction. The bound RNA in the pull-down materials was extracted using TRIzol reagent and analysed using qRT-PCR. Finally, the proteins in the complexes were detected using western blotting. **Table 5** presents the probe sequence used in the study.

### RNA immunoprecipitation

RNA immunoprecipitation (RIP) was conducted using the Magna RIP Kit (Millipore, Billerica, MA, USA) following the manufacturer’s instructions. Antibodies against hnRNPA1, WDR76, Y-box-binding protein 1 (YBX1), and immunoglobulin G used for RIP were purchased from Proteintech Bio. Briefly, the cells were washed two times with PBS and lysed using IP lysis buffer containing RNase and protease inhibitors. The supernatants were collected after centrifugation for further analysis. Next, the antibodies were incubated with protein A/G beads for 1 h at room temperature, and the complexes of antibody and beads were subsequently incubated with cell supernatant overnight at 4°C. RNA in the complex was extracted using TRIzol reagent. Finally, qRT-PCR was used to evaluate the enrichment of circSMAD3 and pre-p53.

### Ethics statement and human aortic tissue acquisition

All mice were maintained and studied using protocols in accordance with the National Institutes of Health Guide for the Care and Use of Laboratory Animals and approved by the Committee on the Ethics of Animal Experiments of the Animal Research Committee of Tongji Medical College, Huazhong University of Science and Technology. Human aortic sections were collected from Tongji Hospital, with approval from the research ethics committees of Tongji Hospital, Tongji Medical College, Huazhong University of Science and Technology.

### Statistical analysis

All statistical analyses were performed using IBM SPSS Statistics for Windows, version 22 (IBM Corp., Armonk, N.Y., USA) and GraphPad Prism 8 (GraphPad Software Inc.). Student’s *t*-test and one- or two-way analysis of variance were used for multiple groups experiment statistical analysis. All data from three independent experiments are presented as the mean±standard deviation (SD) or mean±standard error of the mean (SEM). Statistical significance was set at P < 0.05.

## Results

### CircSMAD3 is the conserved circRNA downregulated in response to vascular injury

We initially performed genomic screening by integrating the circBase database and circRNA microarray data (GSE215935) from mouse aortic tissue to identify the key vascular remodelling-related circRNAs. Additionally, we focused on this gene family to search functional circRNAs since the SMAD family plays a central role in regulating TGF-β1 and is involved in vascular injury^25^. Overall, 34 circRNAs from SMAD1 to SMAD9 genes were identified (Figure 1A). Among them, a previously uncharacterised circRNA, circRNA.19175, which was derived from two to five exons of the SMAD3 gene, known as circSMAD3 (Figure 1B), was upregulated in response to the stimulation of TGF-β1 or serum deprivation and downregulated in response to platelet-derived growth factor subunit B (PDGF-BB) in mASMC (Figure 1C and E). This observation piqued our interest. A conserved human circRNA, hsa_circ_0003973, which contains a homologous junction site for circSMAD3, was also detected in HASMC (Figure 1B). Consistent with the findings of mASMC, hsa_circ_0003973 was largely altered in HASMC after stimulation using TGF-β1, PDGF-BB, and starvation (Figure 1D and F).

**Figure 1.**
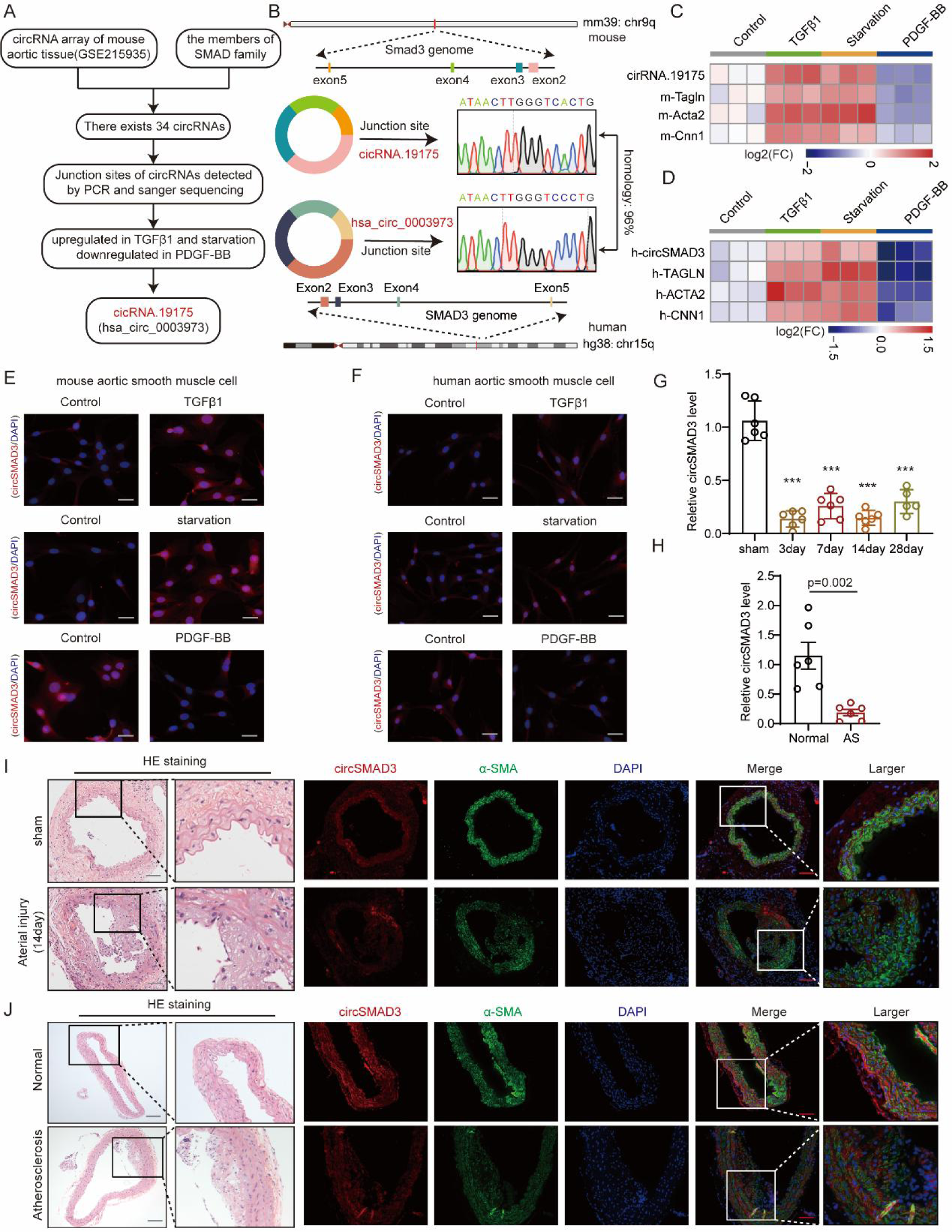
circSMAD3 was significantly downregulated in vascular diseases. **(A)** Schematic identification of circRNAs derived from the members of the SMAD family. **(B)** Schematic of circSMAD3 derived from SMAD3 gene in mouse and human, and their junction sites detected using the Sanger sequencing. **(C-D)** Heat map showing the changes in circSMAD3 in response to TGF-β1 (10 ng/mL), PDGF-BB (20 ng/mL), and starvation for 24 h in mASMC (C) and HASMC (D). **(E-F)** Fluorescence *in situ* hybridisation (FISH) showing the detection of circSMAD3 expression in response to TGF-β1 (10 ng/mL), PDGF-BB (20 ng/mL), and starvation for 24 h in mASMC (E) and HASMC (F). Scale bar=100μm. **(G)** circSMAD3 expression in the time points of arterial injury detected using qPCR. N=6. **(H)** The level of circSMAD3 in atherosclerotic plaque compared with normal aortic tissues via qPCR (N=6). **(I)** qPCR showing the detection of circSMAD3 changes in the injured arterial tissue at different time points. Scale bar=500μm. **(J)** FISH showing the detection of circSMAD3 changes in arterial injury and atherosclerotic plaque. Scale bar=500μm.

In the vascular injury model, circSMAD3 gradually declined in parallel in the carotid artery on days 3, 7, 14 and 28 (Figure 1G and I). Furthermore, circSMAD3 levels were downregulated in mouse atherosclerotic plaques (Figure 1H and J). Therefore, these results indicate that circSMAD3 participated in the pathogenesis of vascular proliferative diseases.

### CircSMAD3 inhibits vascular smooth muscle cell phenotype switching and proliferation *in vitro*

Next, we explored the function of circSMAD3 in phenotype switching and proliferation *in vitro* by infecting mASMCs with sh-circSMAD3 adenovirus and circSMAD3 overexpressing adenovirus, respectively (Figure S1A). CircSMAD3 silencing significantly inhibited phenotype switching by downregulating contractile marker genes, such as α-SMA, smooth muscle protein 22α (SM22α), and Calponin 1 (CNN1), whereas circSMAD3 overexpression dramatically increased these gene expressions (Figure S1B $Figure S2 B-D). Furthermore, circSMAD3 silencing resulted in the switching of VSMCs from elongated contractile morphology to a polygonal synthetic morphology and reversed its contraction effect induced by TGF-β1; however, circSMAD3 overexpression maintained the contractile state of mASMC (Figure S1C-D). Therefore, we performed Transwell, wound healing, EDU, and Cell Counting Kit-8 assays to further investigate the ability of circSMAD3 to affect VSMC proliferation. We found that circSMAD3 silencing substantially promoted the proliferation and migration capacities of VSMCs and receded the effects induced by TGF-β1, whereas its overexpression suppressed these effects (Figure S1E-H). Furthermore, circSMAD3 exerted similar effects on HASMC as the same in mASMCs, and reversed the effects of PDGF-BB in HASMC (Figure 2A-H & Figure S2A).

**Figure 2.**
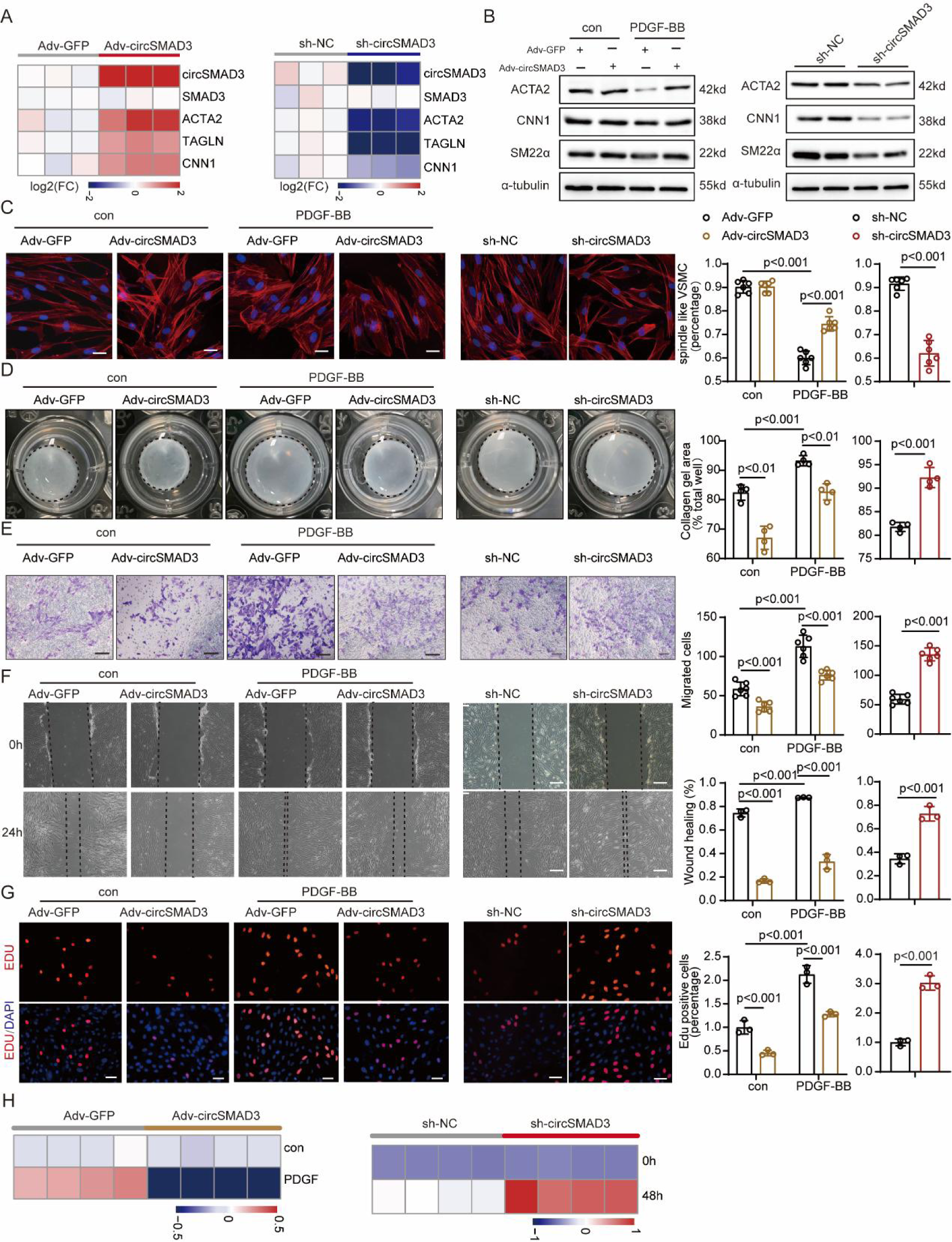
circSMAD3 modulates VSMC phenotype switching and proliferation in HASMC. **(A)** Heat map showing the overexpression and silencing efficiency after being infected with circSMAD3 overexpressing or silencing adenovirus and the expression of contractile marker genes. GAPDH as negative control. Data are shown as mean ± standard deviation (SD) (Student’s t-test, as indicated). **(B)** Western blot showing the detection of the protein level expression of a contractile gene with circSMAD3 overexpression and PDGF-BB (20 ng/mL) (left) and silencing (right). α-Tubulin as the negative control. **(C)** Left, phalloidin staining showing the morphology of HASMC after circSMAD3 overexpression and silencing. Scale bar=50μm. Right, the percentages of spindle-like (contractile phenotype) VSMC in each group. **(D)** Left, collagen gel contrition assay shows HASMC contraction evaluation after circSMAD3 overexpression and silencing. Right, the contraction rate is quantified in each group. Data are shown as mean ± SD (Student’s t-test, as indicated). **(E)** Left, Transwell assay shows HASMC migration evaluation after circSMAD3 overexpression and silencing. Scale bar=200μm. Right, the migration rate is quantified in each group. Data are shown as mean ± SD (Student’s t-test, as indicated). **(F)** Left, wound healing assay to evaluate the migration of HASMC after circSMAD3 overexpression and silence. Scale bar=100μm. Right, the migration rate is quantified in each group. Data are shown as mean ± SD (Student’s t-test, as indicated). **(G)** Left, EDU assay showing the detection of HASMC proliferation after circSMAD3 overexpression and silencing. Scale bar=200μm. Right, the EDU positive rate is quantified in each group. Data are shown as mean ± SD (Student’s t-test, as indicated). **(H)** CCK-8 assay showing the detection of HASMC proliferation after circSMAD3 overexpression and silencing. Data are shown as mean ± SD (two-way ANOVA, as indicated).

### CircSMAD3 attenuates post-injury neointima formation *in vivo*

To assess the role of circSMAD3 in post-injury neointima formation *in vivo*, the carotid arteries of C57BL/6J mice were wire-injured and infected with circSMAD3 overexpression adenovirus at high efficiency (Figure 3A-B). The expression of contractile genes, such as α-SMA, SM22α, and CNN1, were significantly upregulated in the transcriptional and protein levels (Figure 3C). Additionally, Histological analysis and quantification of the arterial neointima/media ratio showed that circSMAD3 overexpression repressed the wire injury-induced neointima formation (Figure 3G-H). Therefore, we evaluated the potential area changes in the neointimal, medial, and luminal regions induced by arterial injury and found that circSMAD3 overexpression prevented the proliferation of VSMCs in the neointima and media regions (Figure 3K). The percentage of Ki67-positive cells representing the proliferation capacity also decreased in the circSMAD3 group (Figure 3K). In contrast, circSMAD3 silencing significantly promoted arterial injury-induced neointima formation (Figure 3D-F & I-J). circSMAD3 knockdown dramatically enhanced the increased proliferation of VSMC induced by injury (Figure 3L). Moreover, contractile genes were markedly downregulated in the circSMAD3 silencing group (Figure 3F).

**Figure 3.**
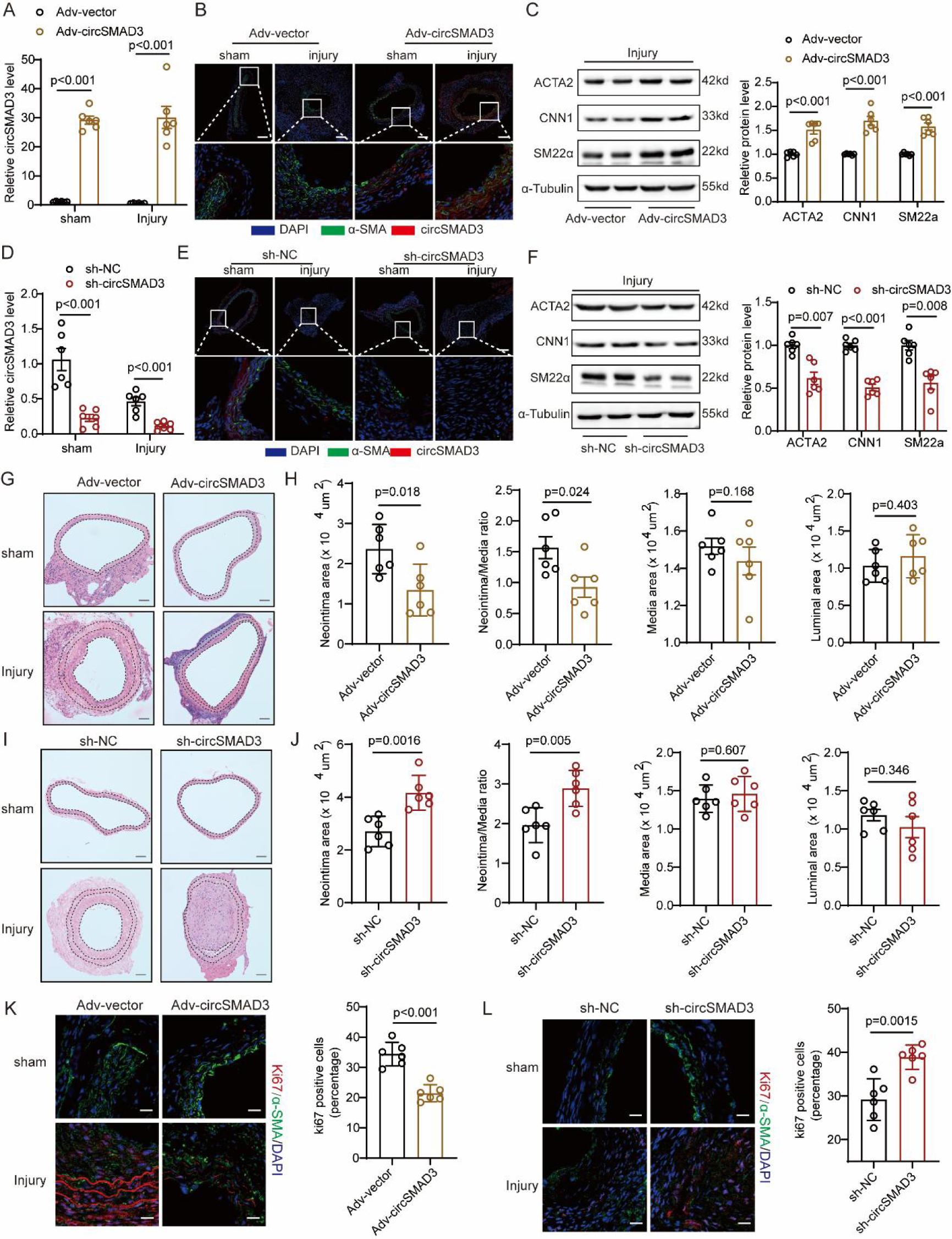
circSMAD3 ameliorated arterial injury-induced neointima formation. **(A&D)** qPCR assays showing the detection of the overexpression efficiency after being infected with circSMAD3 overexpressing adenovirus (A) or sh-RNA adenovirus (D) in carotid arterial tissue. GAPDH as negative control. Data are shown as mean ± standard deviation (SD) (Student’s t-test, as indicated). N=6. **(B&E)** Fluorescence *in situ* hybridisation (FISH) and immunofluorescence showing the detection of circSMAD3 overexpression (B) or sh-RNA adenovirus (E) in carotid arterial tissue. Scale bar=500μm. **(C&F)** Western blot showing the detection of the protein level expression of contractile genes with circSMAD3 overexpression (K) or silencing (L). α-Tubulin as the negative control. Data are shown as mean ± SD (Student’s t-test, as indicated). N=6. **(G&I)** Haematoxylin and eosin (H&E) staining showing the neointima formation caused by arterial injury and circSMAD3 overexpression (C) or silence (F). Scale bar=500μm. **(H&J)** Neointima area, Neointima/media ratio, media area, and luminal area in each group. Data are shown as mean ± SD (Student’s t-test, as indicated). N=6. **(K&L)** Left, Ki-67 staining showing the VSMC proliferation caused by arterial injury and circSMAD3 overexpression. Scale bar=50μm. Right, the percentage of Ki-67 positive cells in each group. Data are shown as mean ± SD (Student’s t-test, as indicated). N=6.

### CircSMAD3 maintains VSMC constriction phenotype through the p53γ signalling pathway

To elucidate the mechanism by which circSMAD3 inhibited VSMC phenotype switching and proliferation *in vitro*, thereby attenuating post-injury neointima formation *in vivo*, we performed transcriptional sequencing to comprehensively describe transcriptomic changes in circSMAD3 silencing HASMC. Gene set enrichment analysis showed that the genes modulated by circSMAD3 were mainly enriched in the DNA replication pathway, p53 signalling pathway, and cell cycle (Figure 4A-C). Since the p53 signalling pathway is critical for cell cycle progression and apoptosis, we hypothesised that circSMAD3 suppresses the proliferation and maintains the contractile state of VSMC by activating the p53 signalling pathway. Therefore, we examined the expression of downstream genes modulated by p53 in response to changes in circSMAD3 expression. Consistent with the transcriptional sequencing results, circSMAD3 silence promoted the expression of cell cycle-related genes (CDK1, CDK2, CCND1, CCNE2, etc) (Figure 4D-E & Figure S3A-B), while the upregulation of cell cycle-related genes induced by PDGF-BB was significantly reduced by circSMAD3 overexpression (Figure 4D-E & Figure S3A-B). Furthermore, when circSMAD3 was knocked down, apoptosis genes, such as FAS and GADD45A, were downregulated, while circSMAD3 overexpression inhibited (Figure 4F &Figure S3C). consistent with the changes of pro-apoptotic genes, the pro-apoptosis effect of circSMAD3 on HASMC was also verified by flow cytometry assay in the condition of circSMAD3 silence or overexpression (Figure 4G & Figure S3D). However, when investigating the role of circSMAD3 in p53, we found that p53 expression was not related to circSMAD3, either at the RNA or protein level, including phosphorylated p53 (Figure 4H-I & Figure S3E-F). What interest us that circSMAD3 knockdown obviously declined the transcriptional function of p53 on its target genes, such as p21, PUMA and NOXA, while overexpression of circSMAD3 enhanced (Figure 4J & Figure S3G). Subsequently, we investigated whether circSMAD3 affected the variants of p53. The results showed that mRNA and p53γ protein were downregulated upon circSMAD3 silencing (Figure 4H-I). However, other variants, such as p53β, Δ40p53, and Δ133p53, had no alterations in response to circSMAD3 changes. circSMAD3 overexpression also promoted the protein level expression of p53γ (Figure S3E-F). Therefore, we conclusively established that circSMAD3 repressed the p53γ signalling pathway, thereby maintaining the differentiated phenotype of VSMCs.

**Figure 4.**
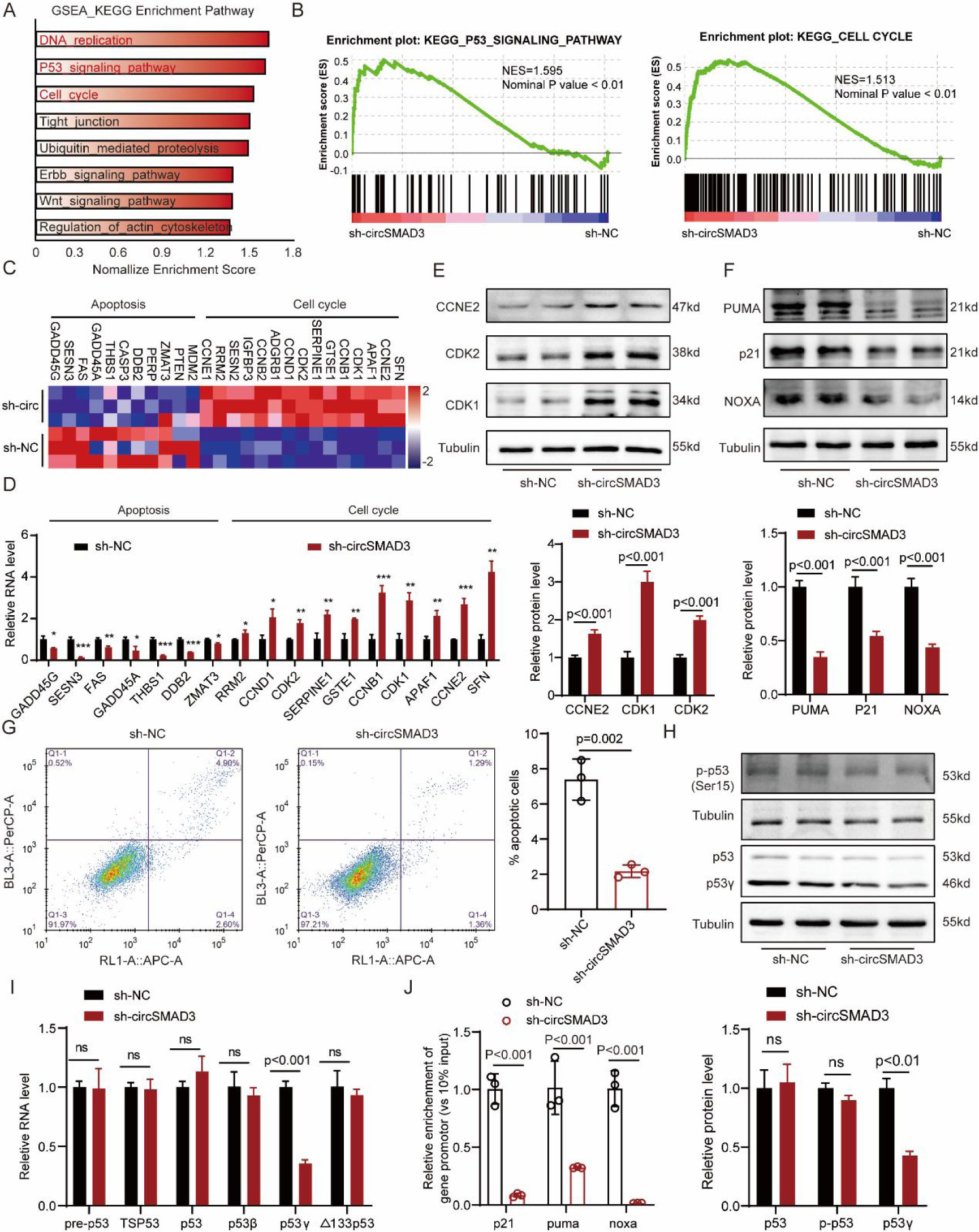
circSMAD3 maintains VSMC phenotype switching through the p53γ signalling pathway. **(A)** The pathways enriched by genes modulated by circSMAD3 silencing were analysed using GSEA. (B) The p53 signalling pathway and cell cycle were the key pathways in response to circSMAD3 silencing. (C) Heatmap showing the apoptosis and cell cycle-related gene expression in the p53 signalling pathway in RNA sequencing. (D) qPCR showing the detection of the changes in apoptosis and cell cycle-related genes in the p53 signalling pathway during circSMAD3 silencing Data are shown as mean ± SD (Student’s t-test, as indicated). (E-F) Western blot to detect proliferative gene expression (E) and pro-apoptosis related genes (F) in circSMAD3 silence. Data are shown as mean ± SD (Student’s t-test, as indicated). (G) Flow cytometry assay to detect the apoptosis of HASMC in circSMAD3 silence. Data are shown as mean ± SD (Student’s t-test, as indicated). (H) Western blot showing the detection of changes in the protein level of p53 isoform and modification level when circSMAD3 is silenced. Data are shown as mean ± SD (Student’s t-test, as indicated). (I) RNA level of p53 isoform in response to circSMAD3 silencing. Data are shown as mean ± SD (Student’s t-test, as indicated). (J) ChIP-qPCR to detect the binding intensity of p53 on its target gene promotors when circSMAD3 silenced. Data are shown as mean ± SD (Student’s t-test, as indicated).

### CircSMAD3 interacts with hnRNPA1 to promote its degradation through the ubiquitination-protease pathway

The interaction of circRNAs with proteins is a classical mechanism by which they perform their function^26^. We identified a set of proteins among the products of circRNA pull-down in HASMC using mass spectrometry. Additionally, we comprehensively collected a set of proteins related to VSMC physiology from the Gene Ontology database and relevant literature and found 11 proteins present in both datasets (Figure 5A). Only hnRNPA1 and YBX1 interacted with circSMAD3 via RNA pull-down and RIP (Figure 5B-C). hnRNPA1, which is an RNA-binding protein, enhances arterial injury-induced neointima formation by modulating pre-RNA splicing^27^. Therefore, we considered hnRNPA1. Bioinformatics analysis using catRapid^28^ and HDOCK^29^ demonstrated that the N-terminal part of hnRNPA1 could bind to circSMAD3 (Figure 5D-E). Therefore, we truncated this protein into three parts with a 3x FLAG tag according to its domains, as shown in Figure 5F. The RNA pull-down assay result showed that circSMAD3 mainly interacted with the RGG-M9 domain of hnRNPA1 (Figure 5F).

**Figure 5.**
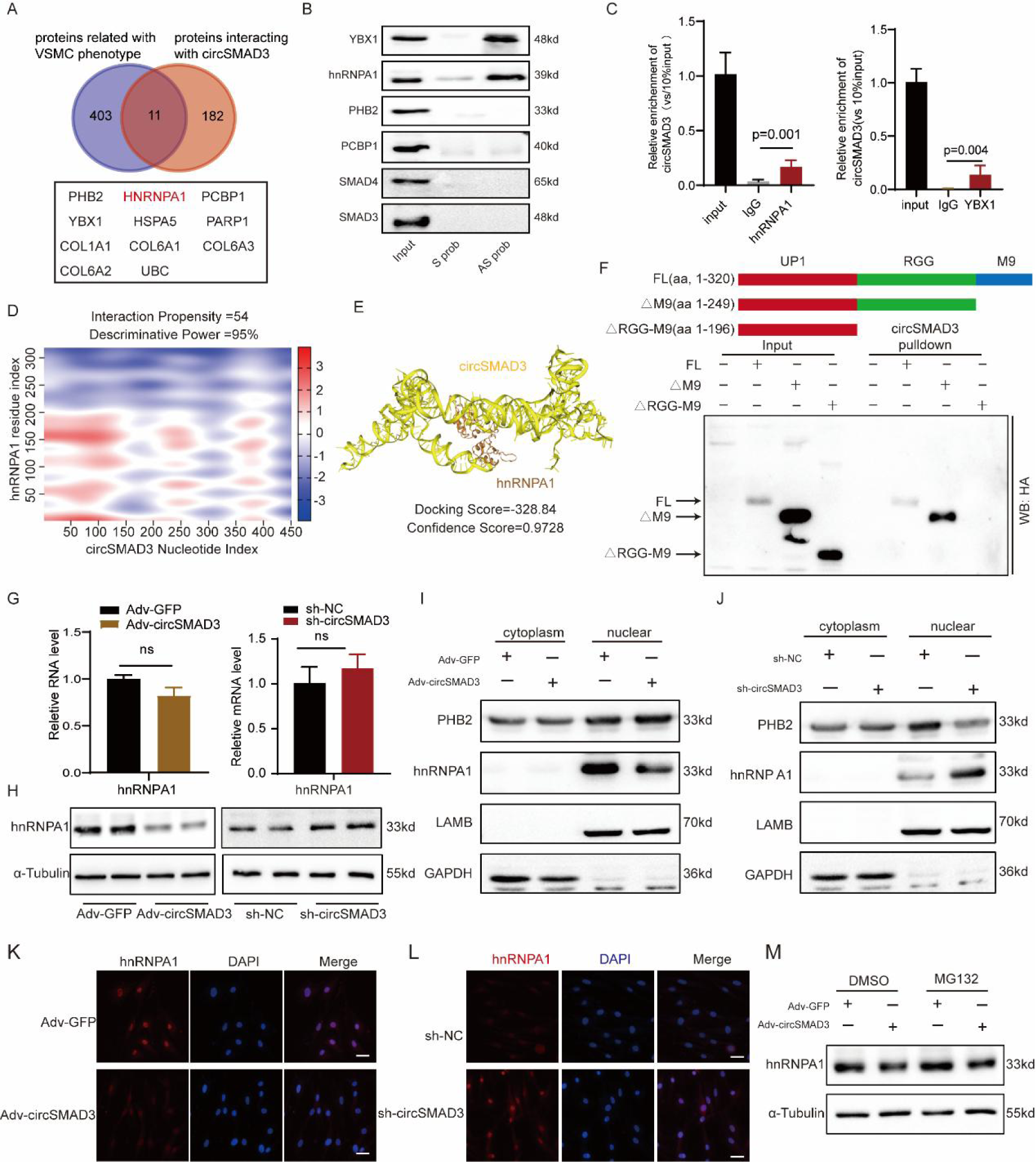
circSMAD3 interacts with hnRNPA1 and promotes its degradation through the protease-ubiquitination pathway. **(A)** The 11 proteins associated with VSMC phenotype and interacting with circSMAD3. **(B)** RNA pull-down to detect circSMAD3 interacted with hnRNPA1 and YBX1. **(C)** RIP assay showing the enrichment of circSMAD3 by hnRNPA1 and YBX1. **(D)** Protein-RNA program-catRapid predicted the N-terminal of hnRNPA1 interacting with circSMAD3. **(E)** HDOCK program for predicting the binding of hnRNPA1 to circSMAD3. **(F)** Truncated assay and RNA pull-down demonstrating circSMAD3 interaction with RGG-M9 region. **(G-H)** The changes in hnRNPA1 in RNA (G) and protein levels (H) when circSMAD3 is overexpressed or silenced. **(I-J)** Cell fraction assay showing the detection of the changes in hnRNPA1 in response to circSMAD3 overexpression (I) and silencing (J) using western blot. **(K-L)** Immunofluorescence assay showing the detection of the changes in hnRNPA1 when circSMAD3 is overexpressed (K) or silenced (L). Scale bar=100μm**. (M)** Western blot showing the detection of the changes in hnRNPA1 protein in response to circSMAD3 overexpression and MG132 (20 ng/mL).

Next, we explored the molecular consequences of the interaction between circSMAD3 and hnRNPA1. CircSMAD3 overexpression and silencing markedly decreased and increased hnRNPA1 protein levels, respectively (Figure 5H). However, we did not observe a reduction in hnRNPA1 transcription (Figure 5G). Reportedly, hnRNPA1 primarily exists in the nucleus to perform its splicing function in pre-RNA. Therefore, we speculated that the influence of circSMAD3 on hnRNPA1 mainly occurs in the nucleus. As expected, cytoplasmic and nuclear fraction separation and immunofluorescence assays showed that when circSMAD3 was overexpressed, the hnRNPA1 protein levels in the nuclear fraction were significantly downregulated, whereas circSMAD3 silencing stabilised hnRNPA1 protein level within the nucleus (Figure 5I-L). Prohibitin 2 (PHB2), which is a protein that reportedly binds with hnRNPA1 to repress its splicing function, was significantly upregulated in response to circSMAD3 overexpression in the nucleus and downregulated when circSMAD3 was silenced (Figure 5I-J). MG132, an inhibitor of the protease-ubiquitination pathway, significantly reversed the circSMAD3-induced reduction in hnRNPA1 protein level (Figure 5M), suggesting that circSMAD3 modulated hnRNPA1 expression, at least partly, through the protease-ubiquitination pathway.

### CircSMAD3 enhances the binding of hnRNPA1 with E3 ligase WDR76

Since WDR76, which is an E3 ubiquitin ligase, has been identified to interact with circSMAD3 through mass spectrometry, we hypothesised that circSMAD3 functions as a scaffold, enhancing hnRNPA1 binding to the E3 ligase WDR76, thereby mediating the degradation of hnRNPA1. In support of this idea, the co-localisation analysis revealed that both hnRNPA1 and WDR76 existed in the nucleus (Figure 6D), and bioinformatic methods predicted a strong interaction between circSMAD3 and WDR76 (Figure 6A-B). Furthermore, RNA pull-down and RIP assays confirmed the binding of circSMAD3 to WDR76 (Figure 6C). As expected, circSMAD3 overexpression enhanced the interaction between hnRNPA1 and WDR76, whereas circSMAD3 silencing abrogated this interaction (Figure 6E-F). Reciprocally, circSMAD3 overexpression facilitated the WDR76-induced ubiquitin level of hnRNPA1 (Figure 6G). Further IP assay showed that WDR76 robustly promoted K63-, K48-, and K33-linked ubiquitination of hnRNPA1 (Figure 6H), which was significantly enhanced by circSMAD3 overexpression (Figure 6I). Therefore, these data suggest that circSMAD3 functions as a scaffold that facilitates the degradation of hnRNPA1 depending on ubiquitination by enhancing the binding of hnRNPA1 to the E3 ubiquitin ligase WDR76.

**Figure 6.**
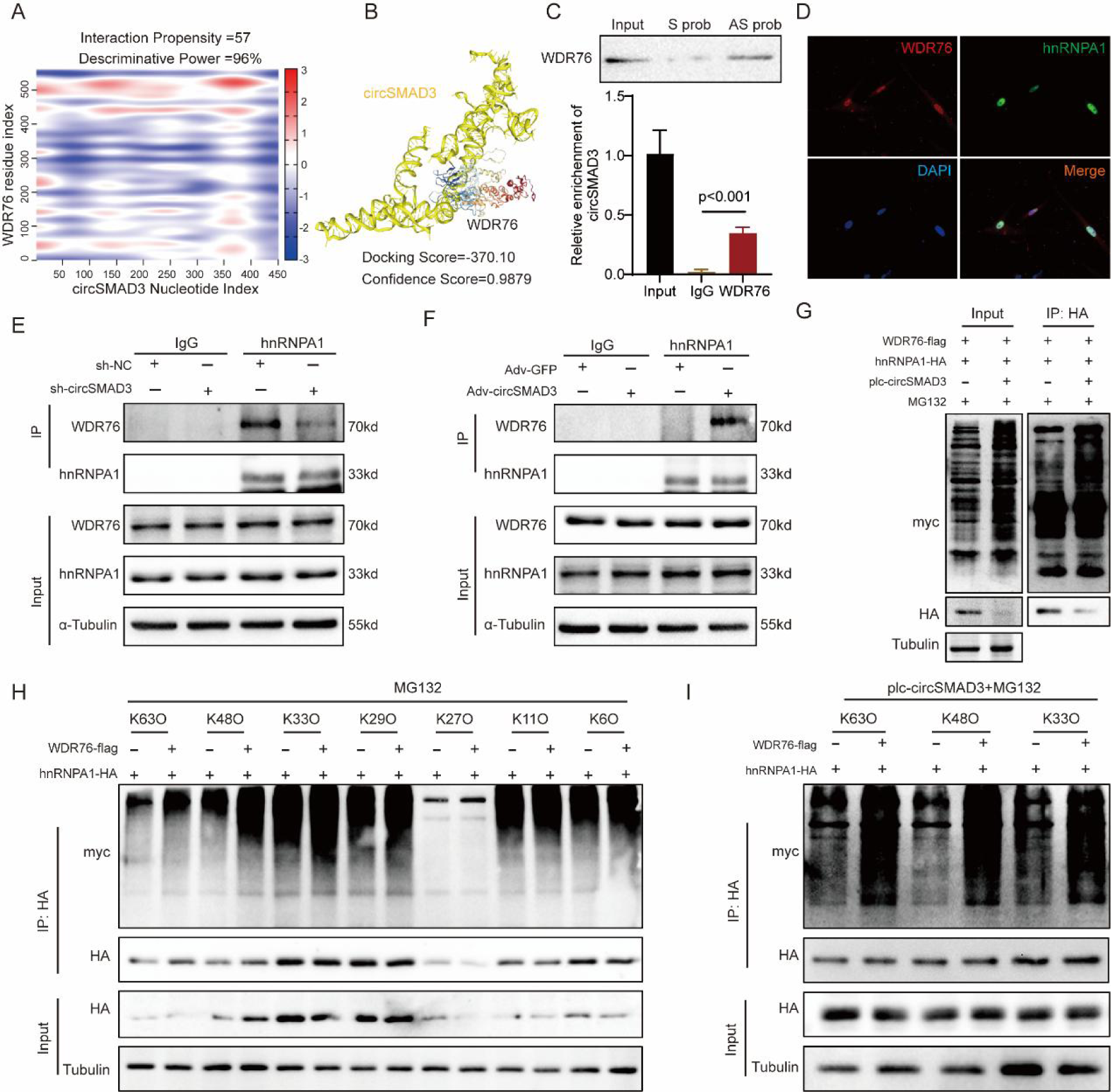
circSMAD3 promoted the ubiquitination degradation of hnRNPA1 by E3 ligase WDR76. **(A-B)** CatRapid and HDOCK predicted that the N-terminus of hnRNPA1 interacts with circSMAD3. **(C)** RIP (upper) and RNA pull-down (lower) showing the detection of the binding of hnRNPA1 to circSMAD3. **(D)** Immunofluorescence assay showing the detection of the co-localisation of hnRNPA1 with WDR76 in HASMC. Scale bar=200μm**. (E-F)** IP assay to detect the interaction between hnRNPA1 and WDR76 upon circSMAD3 silencing (E) and overexpression (F). **(G)** IP assay showing that circSMAD3 overexpression promoted the ubiquitination level of hnRNPA1 in HEK293T cells. **(H)** Western blot showing the ubiquitination levels of hnRNPA1 after Flag-WDR76 overexpression and in response to MG132 treatment in HEK293T cells co-transfected with HA-tagged hnRNPA1 and Myc-tagged ubiquitin (K63O, K48O, K33O, K29O, K27O, K11O, and K6O) constructs. K63O indicates ubiquitin, in which only lysines-K63 were obtained. (H). Western blot showing the ubiquitination levels of hnRNPA1 after circSMAD3 overexpression and in response to MG132 treatment in HEK293T cells co-transfected with HA-tagged hnRNPA1, Flag-WDR76, and Myc-tagged ubiquitin (K63O, K48O, and K33O) constructs.

### CircSMAD3 modulated the alternative splicing of p53 pre-RNA by hnRNPA1

hnRNPA1 is an RNA-binding protein that plays a key role in pre-RNA processing and splicing in the nucleus as a core member of the hnRNP complex^30, 31^. Therefore, we hypothesised that circSMAD3 modulates p53 pre-RNA splicing via the splicing factor hnRNPA1. We found many sites in p53 pre-RNA that interacted with hnRNPA1 using the bioinformatics method catRapid (Figure 7A). Additional evidence revealed that p53 pre-RNA was significantly enriched by hnRNPA1, as detected by the RIP assay, which was significantly promoted by circSMAD3 silencing and decreased by circSMAD3 overexpression (Figure 7B).

**Figure 7.**
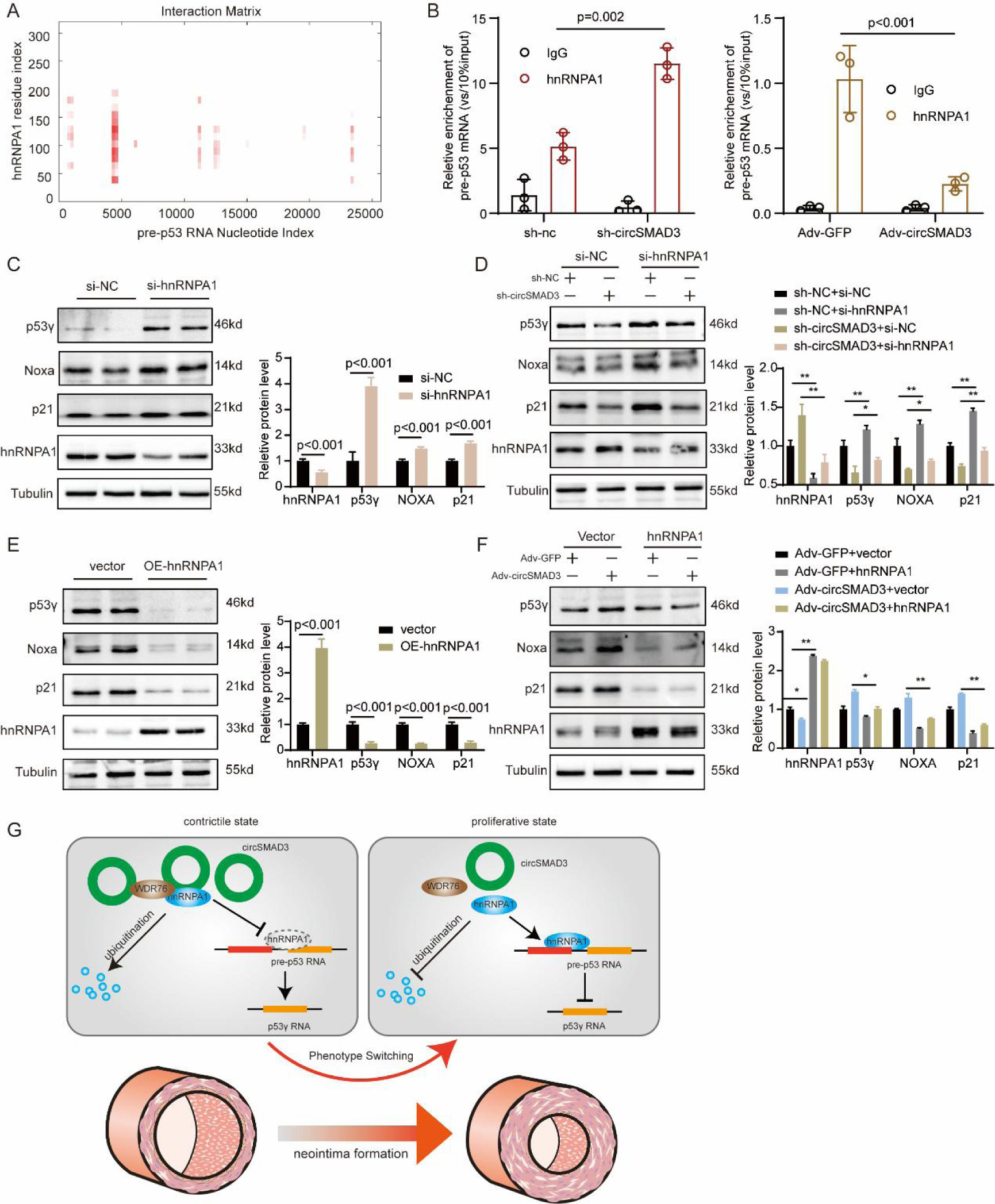
circSMAD3 modulated p53 precursor RNA splicing by hnRNPA1. (A) protein-RNA program-catRapid predicted the binding region of pre-p53 with hnRNPA1. (B) RIP assay showing the enrichment of pre-p53 by hnRNPA1 and the enhancement of this interaction by circSMAD3 silencing and overexpression. (C) hnRNPA1 silencing significantly declined the protein level expression of p53γ ang its target genes. (D) The silencing of hnRNPA1 reversed the downregulation of p53γ ang its target genes induced by circSMAD3 silencing. (E) hnRNPA1 overexpression significantly promoted the protein level expression of p53γ ang its target genes. (F) The silencing of hnRNPA1 reversed the downregulation of p53γ ang its target genes induced by circSMAD3 silencing. (G) Schematic illustration of the mechanism of circSMAD3 in vascular injury. Briefly, when the blood vessel maintains homeostasis, circSMAD3 recruits splicing factor hnRNPA1 to enhance the interaction of hnRNPA1 with E3 ligase WDR76, promote the ubiquitination degradation of hnRNPA1, facilitate p53 pre-RNA splicing to activate the p53γ signalling pathway, and eventually maintain the vascular homeostasis. However, in vascular injury, the downregulated circSMAD3 relieves the repressive function of hnRNPA1 caused by circSMAD3, deactivates the p53γ signalling pathway, and eventually leads to neointima formation of arterial.

Next, we changed hnRNPA1 expression in HASMC to investigate the effect of hnRNPA1 splicing on p53 pre-RNA. hnRNPA1 knockdown promoted the expression of p53γ and apoptosis-related proteins, whereas overexpression of hnRNPA1 decreased (Figure 7C). Additionally, hnRNPA1 silencing abolished the increase in the protein level of p53γ and its target genes induced by circSMAD3 knockdown (Figure 7D). However, hnRNPA1 overexpression rescued the protein level decrease of p53γ and apoptosis-related genes caused by circSMAD3 upregulation (Figure 7E-F). Therefore, these results demonstrated that circSMAD3 modulates the splicing process of p53 pre-RNA mediated by hnRNPA1.

## Discussion

circRNAs have attracted significant attention for developing novel drug therapies because of their safety profile and low immunogenicity. Efficient delivery systems have also accelerated this process significantly. Therefore, elucidating the underlying mechanisms of circRNAs in pathophysiology is essential for developing novel therapeutic targets. Here, we identified a novel circRNA, circSMAD3, which has been shown to significantly affect VSMC proliferation and phenotype switching *in vitro* and *in vivo*. Mechanistically, circSMAD3 interacts with hnRNPA1 to enhance the binding of hnRNPA1 to E3 ligase WDR76, subsequently promoting the ubiquitination degradation of hnRNPA1 to suppress its splicing function on p53 pre-RNA and eventually activating the p53γ signalling pathway (Figure 7G).

An interesting finding of this study is the modulation of circSMAD3 to splice p53 pre-RNA, promote p53γ expression, and enhance the signal transduction of the p53 signalling pathway. In total, 14 isoforms of p53 have been identified because of the alternative splicing of p53 pre-RNA, among which p53β, p53γ, Δ40p53, and Δ133p53 have been extensively explored to play their functions via the p53 signalling pathway^32^. p53β and p53γ enhance the transcriptional modulation of p53 to its target genes by facilitating the binding of p53 to the promotor region; however, this effect is not observed across all p53-inducible promotors^33^. Additionally, Δ40p53 impaired the growth suppression mediated by p53^34, 35^, whereas Δ133p53 inhibited the apoptosis induced by p53^36, 37^. However, the alternative splicing of p53 has rarely been studied in vascular diseases so far, except for circEsyt2 recruiting polyC-binding protein 1 to mediate p53 splicing and facilitate p53β expression^15^. This study found that circSMAD3 participates in the p53 signalling pathway through the transcriptional analysis of HASMC. We also focused on p53, which is a core factor in this pathway. Further investigation confirmed that, even after phosphorylation, circSMAD3 did not alter the transcriptional or post-transcriptional level of p53 expression. We investigated whether the alternative isoforms of p53 participate in the circSMAD3-mediated process. Fortunately, p53γ, rather than other isoforms, had a strong relationship with circSMAD3. Therefore, we speculated that circSMAD3 maintained VSMC homeostasis by activating the p53γ signalling pathway.

Another important finding in this study was the interaction of circSMAD3 with hnRNPA1 to promote its degradation by ubiquitin, which is reported as a splicing factor associated with pre-mRNAs in the nucleus and influences pre-RNA processing. In vascular diseases, hnRNPA1 significantly affects PKM1/2 RNA splicing and reversion from PKM1 to PKM2, ultimately enhancing glycolysis and VSMC phenotype switching. Additionally, we found that hnRNPA1 binds to circSMAD3 by combining the RNA pull-down products in HASMC with proteins related to VSMC physiology. Further truncation experiments confirmed that the RGG-M9 domain in hnRNPA1 was mainly bound to circSMAD3, consistent with PHB2, a direct hnRNPA1 repressor. Additionally, circSMAD3 expression changes synchronously with PHB2 expression in the nucleus, suggesting a synergistic effect between circSMAD3 and PHB2 in modulating VSMC physiology. We also explored the relationship between circSMAD3 and hnRNPA1 and found that circSMAD3 promoted hnRNPA1 degradation via the ubiquitination-protease pathway through the E3 ligase WDR76. Ubiquitination assays also showed that circSMAD3 promoted the K63-, K48-, and K33-linked ubiquitination degradation of hnRNPA1 by enhancing the binding of hnRNPA1 to the E3 ligase WDR76 in HASMC.

We speculated whether hnRNPA1 participates in p53 pre-RNA splicing, considering its role in the alternative splicing of pre-RNA. RIP assay confirmed that the effect of circSMAD3 on p53 pre-RNA splicing was modulated by hnRNPA1. hnRNPA1 silencing in HASMC significantly promoted p53γ expression in the RNA and protein levels, whereas other isoforms had no changes. In contrast, hnRNPA1 overexpression was markedly suppressed. p53γ upregulation induced by circSMAD3 overexpression was immensely abolished by hnRNPA1 overexpression, indicating that hnRNPA1 mediated the process of circSMAD3 modulating p53 pre-RNA splicing.

In this study, we also tried to establish the relationship between circSMAD3 with its parent genes. Unfortunately, circSMAD3 had no effect on SMAD3 expression in transcriptional level and protein level, even its phosphorylated modification. Furthermore, RNA pulldown and RIP assay verified circSMAD3 barely bound with SMAD3. However, circSMAD3 has little effect on the distribution of phosphorylated SMAD3 in cytoplasm and nucleus. But the effect was not enough to attract our attention to further investigate the underlying mechanism.

In conclusion, circSMAD3 suppressed VSMC proliferation and phenotype switching and prevented neointima formation from promoting the interaction of hnRNPA1 with WDR76 to facilitate hnRNPA1 degradation depending on ubiquitination and subsequently activate the p53γ signalling pathway, which may be a novel therapeutic strategy for vascular proliferative diseases.

## Data availability

The RNA-seq data have been uploaded into GEO database and are released in collaboration with NCBI Bioproject (accession number: PRJNA1016164). Additional, RNA-seq raw data will be shared upon reasonable request from the corresponding authors.

## Authors’ contributions

Hu Ding and Jiangtao Yan designed the study and analysed the data. Shuai Mei and Li Zhou conducted the experiments. Shuai Mei, Xiaozhu Ma, Qidamgai Wuyun and Ziyang Cai analysed the data and provided experimental materials. Shuai Mei and Hu Ding drafted the manuscript. All the authors have read and approved this manuscript.

## Supplementary materials

Supplementary information, including the supplementary materials and methods, tables, and figures for this study, is available online.

## Acknowledgements

None.

